# Linking *Gba1* E326K mutation to microglia activation and mild age-dependent dopaminergic Neurodegeneration

**DOI:** 10.1101/2023.09.14.557673

**Authors:** Sin Ho Kweon, Hye Guk Ryu, Hyeonwoo Park, Saebom Lee, Namshik Kim, Seung-Hwan Kwon, Shi-Xun Ma, Sangjune Kim, Han Seok Ko

## Abstract

Mutations in the *GBA1* gene have been identified as a prevalent genetic risk factor for Parkinson’s disease (PD). *GBA1* mutations impair enzymatic activity, leading to lysosomal dysfunction and elevated levels of α-synuclein (α-syn). While most research has primarily focused on GBA1’s role in promoting synucleinopathy, emerging evidence suggests that neuroinflammation may be a key pathogenic alteration caused by GBA1 deficiency. To examine the molecular mechanism underlying GBA1 deficiency-mediated neuroinflammation, we generated *Gba1* E326K knock-in (KI) mice using the CRISPR/Cas9 technology, which is linked to an increased risk of PD and dementia with Lewy bodies (DLB). In the ventral midbrain and hippocampus of 24-month-old *Gba1* E326K KI mice, we found a moderate decline in GBA1 enzymatic activity, a buildup of glucosylceramide, and an increase in microglia density. Furthermore, we observed increased levels of pro-inflammatory cytokines and formation of reactive astrocytes in primary microglia and astrocytes, respectively, cultured from *Gba1* E326K KI mice following treatment with pathologic α-syn preformed fibrils (PFF). Additionally, the gut inoculation of α-syn PFF in *Gba1* E326K KI mice significantly enhanced the accumulation of Lewy bodies in the dentate gyrus of the hippocampus, accompanied by aggravated neuroinflammation and exacerbated non-motor symptoms. This research significantly enhances our understanding of the *Gba1* E326K mutation’s involvement in neuroinflammation and the cell-to-cell transmission of pathogenic α-syn in the brain, thereby opening new therapeutic avenues.

## Introduction

Mutations in *GBA1* gene are recognized as the common genetic risk factor for developing Parkinson’s disease (PD)^1^. Glucocerebrosidase (GCase; EC 3.2.1.45, encoded by *GBA1* gene) is a lysosomal hydrolase involved in glucocerebrosidase metabolism^2^. Hetero- and homozygous *GBA1* mutations reduce its enzymatic activity, which may result in lysosomal dysfunction and increased α-syn levels^3^. While most studies focus on the role of GBA1 in promoting synucleinopathy, neuroinflammation may be a key pathogenic alteration caused by GBA1 loss. Neurodegenerative diseases like Alzheimer’s disease (AD), PD, and dementia with Lewy bodies (DLB) are linked to neuroinflammation. This response also is characterized by the activation and proliferation of microglia in the brains of individuals affected by type 2 Gaucher disease (GD)^4^. PD patients with *GBA1* mutations exhibit increased levels of inflammatory markers and cytokines in their plasma, including IL-8, monocyte chemotactic protein 1, and macrophage inflammatory protein 1α^5^. Accumulation of glucosylceramide (GlcCer), a substrate for GCase, may activate complement systems and aggravate immunological complexes, resulting in unintended release of pro-inflammatory cytokines and chemokines^6^. Notably, the systemic inhibition of GBA1 by CBE (Conduritol B epoxide) treatment results in the upregulation of complement C1q^7^ and an increase in Receptor-Interacting Protein Kinase 3 (RIPK3) levels in microglia of layer V of the cortex^8^. Massive activation of microglia has been observed in the mouse model of CNS-restricted GBA1 deficiency^9,10^. Furthermore, both GD and PD patients with *GBA1* mutations have astrogliosis and aberrant α-syn inclusion^11^. Consistent with this observation, iPSC-derived astrocytes harboring *GBA1* mutations also exhibit high proliferating property with overlapping astrocytes processes and secretion of MCP1, CXCL1 and IL-8 in response to monomeric α-syn^12^.

Recently, *GBA1* E326K variant, which has been observed in individuals with Parkinson’s disease (PD) but not Gaucher disease (GD), has emerged as a significant risk factor for DLB^1,13–16^. PD patients carrying a *GBA1* E326K variant have been specifically associated with a distinct pattern of cognitive impairments in working memory, executive function, and visuospatial abilities^15,17^. The GBA1 E326K protein shows a characteristic ER-retention pattern and reduced trafficking to the lysosome^18^. GBA1 deficiency leads to lysosomal accumulation of its lipid substrates, such as GlcCer and glucosylsphingosine (GlcSph), resulting in impaired lysosomal function. This includes alterations in lysosomal content and an elevation in lysosomal pH^19–22^. However, it remains unclear whether the deficiency of GBA1 due to the E326K mutation is involved in neuroinflammation that contributes to neurodegeneration in PD.

In the present study, we generated *Gba1* E326K knock-in (KI) mice using the CRISPR/Cas9 technology to examine the molecular mechanisms underlying GBA1 deficiency-mediated neuroinflammation. We report that the homozygous *Gba1* E326K mutation led to a reduction in both the levels and activity of GCase, accompanied by an accumulation of GlcCer. Moreover, we observed increased neuroinflammation and neuronal loss in the hippocampus of *Gba1* E326K KI mice. Additionally, we observed exacerbated activation of microglia and the formation of reactive astrocytes in α-synuclein (α-syn) preformed fibril (PFF)-induced *Gba1* E326K primary glial cells. Notably, when α-syn PFF was administered via gut injection, it induced exacerbated α-syn pathology and neuroinflammation in the hippocampus of *Gba1* E326K KI mice. Furthermore, the gut transmission of pathogenic α-syn PFF worsened non-motor behavioral deficits in *Gba1* E326K KI mice. These findings strongly support the link between the *Gba1* E326K mutation, neuroinflammation, α-synuclein pathology, and neurodegeneration in PD.

## Materials and Methods

### Mice

All animal experiments were according to the guidelines of Laboratory Animal Manual of the National Institute of Health Guide to the Care and Use of Animals and were approved by the Johns Hopkins Medical Institute Animal Care and Use Committee. We have taken significant steps to alleviate animal pain and discomfort. *Gba1* E326K conditional knock-in (KI) mice were generated by homologous recombination using a targeting vector containing exon 8, which is flanked by loxP sites followed mutated exon 8 (g1030G>A, Glu326Lys), and two sgRNA (1st sgRNA: 5’-gggccctggaaagttcagagtgg-3’, 2nd sgRNA: 5’-tgtgaaagagaagataaccatgg-3’). Due to neonatal lethality of floxed mice, all *Gba1* E326K KI mice used to this study were obtained after mating with CMV-Cre^+^ mice to remove flanked region by loxP sites. CMV-Cre^+^ mice were obtained from the Jackson Laboratories (ME, USA). The mouse tail was treated with 200 μl of DirectPCR Lysis reagent (Viagen Biotech, CA, USA) with 0.5 mg/ml of Proteinase K (Sigma-Aldrich, MO, USA) at 55°C for 24 h. Subsequently, genomic DNA was isolated for genotyping by incubation at 85°C for 45 min. 2μl of the crude lysates was added to PCR reaction buffer containing 10 mM dNTPs, 10 μM forward and reverse primers, 1 μl of AmpliTaq® DNA polymerase with 1X GeneAmp® PCR buffer II (Applied Biosystems). PCR reaction was conducted using Veriti thermal cycler (Applied Biosystems), with the following PCR conditions: 1) 95°C for 5 min, 35 cycles of 95°C for 30 sec, 51.5°C for 30 sec, 72°C for 30 sec, followed 72°C for 7 min. The primer sequences used for the PCR were as follows: forward, 5’-ACT CTG AAC TAT AAC TTC GT-3’; reverse, 5’-AAA GAC CGA TGG CCA TAC AC-3’, PCR for the wild type was no product. For KI, PCR product size was approximately 521 bp. 2) 95°C for 4 min, 35 cycles of 95°C for 1 min, 61°C for 15 sec, 72°C for 1 min, followed 72°C for 7 min. The primer sequences used for PCR were as follow: forward, 5’-TGT GTG GGC CTT ATC TGA CA-3’; reverse, 5’-AGG GCC CTG GAA AGT TCA-3’, PCR product size for the wild type was approximately 309 bp. PCR for KI was no product. The PCR product were electrophoresed on 2% agarose gel and determined the genotype of mice.

### Lysosomal enriched fraction and measurement of lysosomal GCase activity

The enzymatic assays were conducted using the lysosomal-enriched fraction as previously described method^23^. In short, the lysosome-enriched pellet was then retrieved in 50 μl of GCase activity assay buffer (0.25% Taurocholic acid, 1 mM EDTA, 0.25% Triton X-100, in citrate/phosphate buffer, pH 5.4). Samples were resuspended in 50 μl of 1 mM 4-Methylumbelliferyl b-glucopyranoside (4-MU) with or without 10 mM conduritol B epoxide with 1% BSA to measure the GCase activity. After stopping the reaction with the stop solution (50 μl of 1 M glycine, pH 12.5), the fluorescence was measured with a Perkin Elmer plate reader (ex = 355 nm, em = 460 nm, 0.1 s). Lysates in lysosomal-enriched fraction was also subjected to immunoblot analysis.

### Immunoblot analysis

Using a Diax 900 homogenizer (Sigma-Aldrich), mouse brain tissues were homogenized in lysis buffer (10 mM Tris-HCl, pH 7.4, 150 mM NaCl, 5 mM EDTA, 0.5% Nonidet P-40, 10 mM Na-β-glycerophosphate) containing phosphate inhibitor mixture I and II (Sigma-Aldrich) and complete protease inhibitor mixture (Roche, IN, USA). Following homogenization, samples underwent an incubation time of 30 min at 4°C to achieve full lysis. To obtain the supernatants, these lysates were centrifuged at 22,000 g for 20 min. For Triton X-100 soluble and insoluble fraction, cells were prepared using sequential lysis buffer. The following Triton X-100 soluble buffer was used to homogenize the samples in buffer containing 50 mM Tris (pH 8.0), 150 mM NaCl, 1% Triton X-100 with a protease inhibitor cocktail. After centrifugation, the soluble supernatant was collected. The insoluble pellet was resuspended in a solution of 50 mM Tris (pH 8.0), 150 mM NaCl, 1% Triton X-100, 2% SDS, and a protease inhibitor cocktail with sonication (10% amplitude, 1 sec pulse on/off, total 15 sec), centrifuged at 22,000 x g for 30 min, and then supernatant was collected. The BCA assay (Pierce, Rockford, IL, USA) was used to measure protein concentrations. Lysates were prepared by utilizing 2 × Laemmli buffer (Bio-Rad). Proteins from mouse brain tissue were separated on 8%–16% and 4%–20% gradient SDS-PAGE gels before being transferred to nitrocellulose membranes. After blocking the membrane with a solution containing Tris-buffered saline with 5% non-fat dry milk and 0.1% Tween-20 for 1 h, the membrane was then incubated at 4°C overnight with anti-LAMP1 (1:1,000, Cell Signaling), anti-GBA1 (1:1,000, Sigma-Aldrich), anti-α-synuclein (1:1,000, BD Biosciences), anti-pSer129-α-syn (1:30,000, Cell Signaling), anti-Vimentin antibodies (1:2,000, Cell Signaling), anti-C3 (1:1,000, Abcam) antibodies, anti-m-CSFR (1:1,000, Cell Signaling), and anti-GFAP (1:1,000, Dako), followed by 1 h at room temperature with HRP-conjugated rabbit or mouse secondary antibodies (1:50,000, GE Healthcare). Enhanced chemiluminescence (Thermo Scientific, IL, USA) was used to see the bands. After a previous antibody was stripped, the membranes were examined once again using an HRP-conjugated β-actin antibody (1:40,000, Sigma-Aldrich).

### Filter-trap assay

Samples were placed into the pre-wetted nitrocellulose membrane in the Bio-Dot microfiltration apparatus (Bio-rad). Under vacuum, 20 μg of tissue lysates were loaded onto the membrane. Following a Tris-buffered saline wash, samples were blocked in a solution that contained 0.1% Tween-20 and 5% non-fat dry milk. Membranes were incubated with anti-A11 oligomers antibody (1:500, Life technologies) at 4°C overnight, followed by HRP-conjugated rabbit secondary antibody (1:50,000, GE Healthcare) for 1 h at room temperature and the signals were visualized using chemiluminescence reagents (Thermo Scientific).

### Immunohistochemistry

Mice were fixed with 4% paraformaldehyde/PBS (pH 7.4) after receiving an ice-cold PBS perfusion. After being collected, the brains were post-fixed for 16 h in 4% paraformaldehyde and followed by cryoprotected in a solution of 30% sucrose/PBS (pH 7.4) for 3 days. An optimal cutting temperature (OCT) buffer was used to freeze the brains before cutting 30 µm serial coronal pieces with a microtome. Free-floating 30 µm sections were incubated with an antibody against TH (1:1,000, Novus Biologicals; 1:1,000, Sigma-Aldrich) or an antibody against Iba1 (1:1,000, Wako) followed by incubation with a biotin-conjugated anti-rabbit or anti-mouse antibody (1:250, Vectastain Elite ABC kit, Vector laboratories). Sections were counterstained with Nissl (0.09% thionin) after being developed using SigmaFast DAB Peroxidase Substrate (Sigma-Aldrich). With the use of a computer-assisted image analysis system composed of an Axiophot photomicroscope (Carl Zeiss Vision) fitted with a computer controlled motorized stage (Ludl Electronics), a Hitachi HV C20 camera, and Stereo Investigator software (MicroBright-Field), TH-positive and Nissl-positive DA neurons from the SNc region were counted through optical fractionators, which is the unbiased method for cell counting. The number of Iba1-positive microglia in the SNc and hippocampus regions was measured with ImageJ software. Four areas of interest within each tissue section were analyzed to estimate the average number of Iba1-positive microglia by field counting, and the data is presented as Iba1 positive cells per mm^2^.

### Immunofluorescent analysis

4% Paraformaldehyde/PBS (pH 7.4)-fixed 30 µm coronal brain sections in SNc, hippocampus, and amygdala regions were blocked with 10% donkey serum (Jackson Immunoresearch)/PBS plus 0.1% Triton X-100 and incubated with antibodies to TH (1:1,000, Novus Biologicals) and/or GlcCer (1:500, Cedarlane), pSer129-α-syn (1:1,000, Biolegend), NeuN (1:1,000, Millipore), Iba1 (1:1,000, Wako) and GFAP (1:1,000, Dako) overnight at 4°C. Following PBS washes, floating brain sections were incubated with 0.1% Triton X-100 and 5% donkey serum in PBS. Thereafter, at room temperature, they were incubated for 1 h with a combination of FITC- and CY3-conjugated secondary antibodies (1:250, Jackson Immunoresearch). After mounting the coverslips with DAPI mounting medium (VECTASHIELD HardSet Antifade Mounting Medium with DAPI, Vector laboratories), the fluorescence images were captured using a Zeiss confocal microscope (Zeiss Confocal LSM 710). The Zeiss Zen software was used to process each image. ImageJ analysis was used to calculate the size of the chosen region inside the threshold’s signal intensity range.

### Primary glia and cortical neuron culture

Primary microglia and astrocyte were obtained from the brains of postnatal mouse pups (P1-3). The extracted brains were washed in Hanks’ Balanced Salt solution (HBSS) three times, and then brains were treated with 0.25% trypsin–EDTA (Gibco) and removal of cell debris and aggregates with a 100 μm nylon mesh for single cell suspension. The single cell suspension was cultured with DMEM/F12 (Gibco) supplemented with 10% heat-inactivated FBS, 50 U/ml penicillin, 50 μg/ml streptomycin, 2 mM L-glutamine, 100 μM non-essential amino acids, and 2 mM sodium pyruvate in T-175 flasks for 21 days, with a complete medium change every 7 days. The mixed glial cell cultures were separated into a magnetically bound microglia-enriched fraction and pour-off astrocyte-enriched fractions using the EasySep Mouse CD11b Positive Selection Kit (StemCell technologies) in accordance with the manufacturer’s instructions. After 24 h, microglia and astrocyte were kept in FBS-free DMEM/F12 supplemented with 50 U/ml penicillin, 50 μg/ml streptomycin, 2 mM L-glutamine, 100 μM non-essential amino acids, and 2 mM sodium pyruvate. After 4 h, microglia were incubated with 5 μg/ml of α-syn PFF for 24 h, and then collected cells and microglia-conditioned medium (MCM). Astrocyte were incubated with MCM for 24 h, and then collected cells and astrocyte-conditioned medium (ACM).

Cortical neurons were prepared from embryonic day 14.5 from pregnant C57BL/6 mice (Charles River). Primary cortical neurons were cultured in Neurobasal media supplemented with B-27 (Gibco), 0.5 mM L-glutamine (Gibco), penicillin and streptomycin (Invitrogen) on tissue culture well plates coated with 100 μg/ml poly-_D_-lysine (Sigma-Aldrich). Neurons were added fresh media every three days. On days *in vitro* (DIV) 14, neurons were treated with 50 μg/ml of ACM for 36 h, followed by cell viability assay using the AlamarBlue reagents (Invitrogen). After 4 h incubation of AlamarBlue solution, cell viability was determined by fluorescence using a Perkin Elmer plate reader (ex = 570 nm, em = 585 nm, 0.1 s).

### RT-PCR

Total RNA was isolated from primary microglia or astrocyte using the RNeasy Plus Mini kit (Qiagen). The cDNA was subsequently synthesized with an Invitrogen SuperScript® IV First-Strand Synthesis System. The relative levels of mRNA expression were examined utilizing real-time PCR (Applied Biosystems ViiA 7 Real-Time PCR System) with SYBR® GreenER™ reagent (Invitrogen) in accordance with the manufacturer’s instructions. The 2−ΔΔCT method was used for calculating the values. All ΔCT values were normalized to Actin. Primer sequences for RT-PCR are summarized in supplementary Table 1.

### PFF generation

The mouse recombinant full-length α-Syn monomer was purified by following the instructions^24^. Briefly, after serial purification steps using Superdex 200 Increase 10/300 G size-exclusion and Hitrap Q Sepharose Fast Flow anion-exchange columns (Cytiva), the α-Syn monomers were subjected to HiTrap SP HP cation-exchange column (Cytiva) for removing lipopolysaccharides. The α-Syn fibrils were mixed together by stirring in an Eppendorf orbital mixer for 7 days at 1,000 rpm and 37°C. Before use, the α-Syn monomer was kept at 80°C. α-Syn fibrils were sonicated for 30 sec (0.5 sec pulse on/off) using a Branson Digital sonifier with a 10% amplitude.

### Gastrointestinal injection of PFF

PFF was injected into the gastrointestinal tract as previously mentioned^24^. Briefly, 3 months old mice were anesthetized with isoflurane (2%–4%) and given 2 injections into the pyloric stomach wall and 2 injections into the duodenal intestine wall, each 0.5 cm apart from the pyloric stomach injection site. The upper duodenum received 6.25 μg of α-Syn PFF in two distinct places (2.5 μg/μl, total 2.5 μl/location) for a total of 12.5 μg, while the pyloric stomach received 6.25 μg of syn PFF in two different areas (2.5 μg/μl, total 2.5 μl/location) for a total of 12.5 μg. Additionally, control mice received the same amounts of PBS injections in the same places.

### Behavior tests

#### Pole test

The behavioral process room was used to acclimate the mice for 30 min. The pole was made of a 9 mm-diameter, 75 cm-long metal rod. The pole was covered in bandage gauze. Mice were positioned on top of the pole with their heads up and 7.5 cm below the top of the pole. The total amount of time it took to get to the pole’s base was noted. Mice underwent two consecutive days of training before the test day. In each training session, three test trials were conducted. Mice were tested three times on the test day, and the total testing time was noted. The test and recording could only be stopped for a total of 60 sec. Results were recorded for the climb down, turn down, and overall time.

#### Grip strength

Neuromuscular strength testing was carried out utilizing a Bioseb grip strength test device (BIO-GS3, Bioseb). The mice’s performance was evaluated two times. Mice were permitted to hold a metal grid with their forelimbs in order to measure their grip strength. When the mice relinquished their hold on the grid, the force transducer measured the tail’s maximum holding force. Digitally, the maximum holding force was captured and shown as force in grams.

#### Morris water maze test

The Morris water maze test has four separate inner cues on the surface and is a white, round pool that is 150 cm in diameter and 50 cm high. The platform was submerged 1 cm below the water’s surface so that it was invisible at water level. The circular pool was filled with water and a nontoxic water-soluble white dye (20 ± 1°C). The swimming pool was divided into four equal-sized quadrants. Within one of the pool’s four quadrants, a black platform (9 cm in diameter and 15 cm high) was positioned in the middle. Using a video tracking system (ANY-Maze), the position of each swimming mouse from its starting point to the platform was digitally recorded. Mice underwent swim training for 60 seconds without the platform the day before the trial. The mice were subsequently subjected to two trial sessions per day for four days in a row, with a 15-min intertrial gap. The escape latencies were then recorded. For each trial session and each mouse, this parameter was averaged. The mouse was given permission to stay on the platform for 10 sec after finding it. If the mouse failed to locate the platform in 60 sec, it was put there for 10 sec before being put back in its cage by the experimenter. On day 6, mice were given a cut-off time of 60 sec for the probe trial test that involved removing the platform from the water.

#### Step-through passive avoidance test

The step-through passive avoidance device has a guillotine door separating the clear chamber from the dark chamber. 2 mm stainless steel rods spaced 0.5 cm apart made up the floor of the clear chamber (36 cm x 18 cm x 30 cm) and the dark chamber (36 cm x 18 cm x 30 cm) (Coulbourn Instruments, Holliston, MA, USA). The equipment was lighted by the cue light that was placed above the clear chamber. The mice underwent two separate trials: a training trial and a test trial 24 h later. Mice were initially placed in the clear chamber for the acquisition trial. The door closed as mice entered the dim box and the stainless steel rods delivered a 0.5 mA, 3-sec electrical foot shock. Mice were moved back into the lit compartment for the retention trial 24 h after the retention trial. For both training and test trials, latency was the length of time it took for a mouse to enter the dark compartment after the door opened. It took up to 300 sec for latency to reach the dark compartment.

#### Y-maze test

A spontaneous alternation behavior Y-maze test consists of a horizontal maze with three equal angles between each arm that is 40 cm long, 10 cm broad, and has walls that are 15 cm high. The walls and floor of the maze were built from opaque polyvinyl plastic. Mice were originally placed within one arm, and over the course of 8-min, the order and quantity of arm entrances were manually recorded for each mouse. Precision short-term memory was recorded as a spontaneous alternation when the mice entered all three arms. The ratio of the actual alternations to the possible alternations (defined as the total number of arm entries minus two) was multiplied by 100 to determine the alternation score (%) for each mouse: [(Number of alternations)/(Total arm entries – 2)] ×100. Locomotor activity was measured by the number of arm entries per trial. In order to remove smells and contaminants between experiments, the Y-maze arms were cleaned with 10% ethanol that had been diluted.

#### Novel object recognition test

A grey open field box with opaque polyvinyl plastic (45 cm in width, 45 cm in depth, and 50 cm in height) was used for novel object recognition test. All mice were introduced to the test box for 5 min without any objects before the test. Mice were placed into the open field box with two identical objects after the introduction phase and given 5 min to explore. Wooden blocks that were the same size but had a varied form were the objects utilized in this experiment. We recorded how long the mice spent investigating each object during the training period. Next day mice were allowed to search the objects for 5 min, at which time a familiar object from the training was placed next to a novel object. The length of time the mice spent investigating the novel and the recognizable objects was noted. Following each session, 10% ethanol was used to clean the items and test box. The percentage of time spent recognizing novel objects is used to express the data (time percentage = total time spent with novel object/[total time spent with novel object + total time spent with familiar object] × 100).

#### Open-field test

A rectangular plastic box (40 cm x 40 cm x 40 cm) divided into 36 identical sectors (6.6 cm x 6.6 cm) made up the open field. The field was divided into two sectors: the core sector, which contained four center squares (2 by 2), and the periphery sector, which contained the remaining squares. The mouse was left in the center of an open field for 5 min in dim light to explore. Between each trial, the device was carefully cleaned using 10% diluted ethanol. The distance traveled was recorded using a video tracking system (ANY-Maze software) as a gauge of locomotor activity. As a measure of anxiolytic activity, the length of stay and number of visits to the center were recorded.

#### Elevated plus maze

The elevated plus maze had two open arms (50 cm x 10 cm) and two closed arms, each of which had a wall (30 cm x 5 cm x 15 cm) connecting them to the center zone (5 cm x 5 cm), making a cross. It had been raised 50 cm above the ground. To record the experiment, a camera was hung from above the maze. The open arms had a modest (0.5 cm) edge, and the walls and floor of the maze were made of opaque polyvinyl plastic. Each mouse was positioned at the center junction with its back to an open arm. To get rid of smells and contaminants, the maze floor was properly cleaned in between testing using 10% diluted ethanol. Using the Any-Maze behavioral tracking system, the length of time spent in the open arms and the number of entrances during a 5-min period were both recorded. An entry was defined as all four paws in the arm. The length of time in the open arms and the quantity of entries into the open arms were transformed, respectively, into percentages of the total time and entries.

#### Forced swimming test

Mice were briefly exposed to a glass cylinder (20 cm in height, 14 cm in diameter) filled 16 cm-high with water (25 ± 1° C). Following a 6-min forced swim test, the video tracking system (ANY-Maze software) was used to calculate and analyze the immobility time during the test’s final 4-min interval. Time spent immobile was compared to a mouse passively floating on the water, making only little movements to keep its snout above the water.

#### Tail suspension test

Each mouse was separately suspended in black Plexiglas boxes (50 cm x 50 cm x 50 cm) by an adhesive tape that was positioned about 1 cm from the tip of the tail and hung 5 cm above the ground. During a 6-min test, the immobility time was noted using the video monitoring system (ANY-Maze program).

### Statistics

Data from at least 3 separate experiments were given as mean ± SEM. An unpaired two-tailed Student’s test or one- or two-way ANOVA test followed by Bonferroni post-hoc multiple comparison analysis was performed to assess the statistical significance using GraphPad Prism 9 software. Assessments with p < 0.05 were considered significant.

## Results

### The homozygous mutation *Gba1* E326K leads to a reduction in both the levels and activity of GCase, which is accompanied by neurodegeneration

To determine the effect of the *Gba1* E326K mutation in a mouse model, we employed the CRISPR-Cas9 and Cre-loxP systems to generate inducible *Gba1* E326K knock-in mice (Supplementary Fig. S1A). The generation of *Gba1* E326K knock-in mice involved the simultaneous injection of two sets of gRNAs and donor DNA containing LoxP targeted to the *Gba1* locus, followed by the breeding of the CMV-Cre-driver strain with a floxed mouse strain (Supplementary Fig. S1B and C). First, we explored the effect of the *Gba1* E326K heterozygous and homozygous mutations on the activity and protein levels of brain glucocerebrosidase (GCase) in hippocampal tissues collected from *Gba1*^WT/WT^, *Gba1*^+/E326K^, and *Gba1*^E326K/E326K^ mice at 24 months of age. As described previously^25,26^, the GCase activity (Fig. 1A) and protein levels (Fig. 1B and C) in the hippocampus of *Gba1*^E326K/E326K^ mice were reduced by approximately 50% compared to those in WT mice.

**Figure 1.**
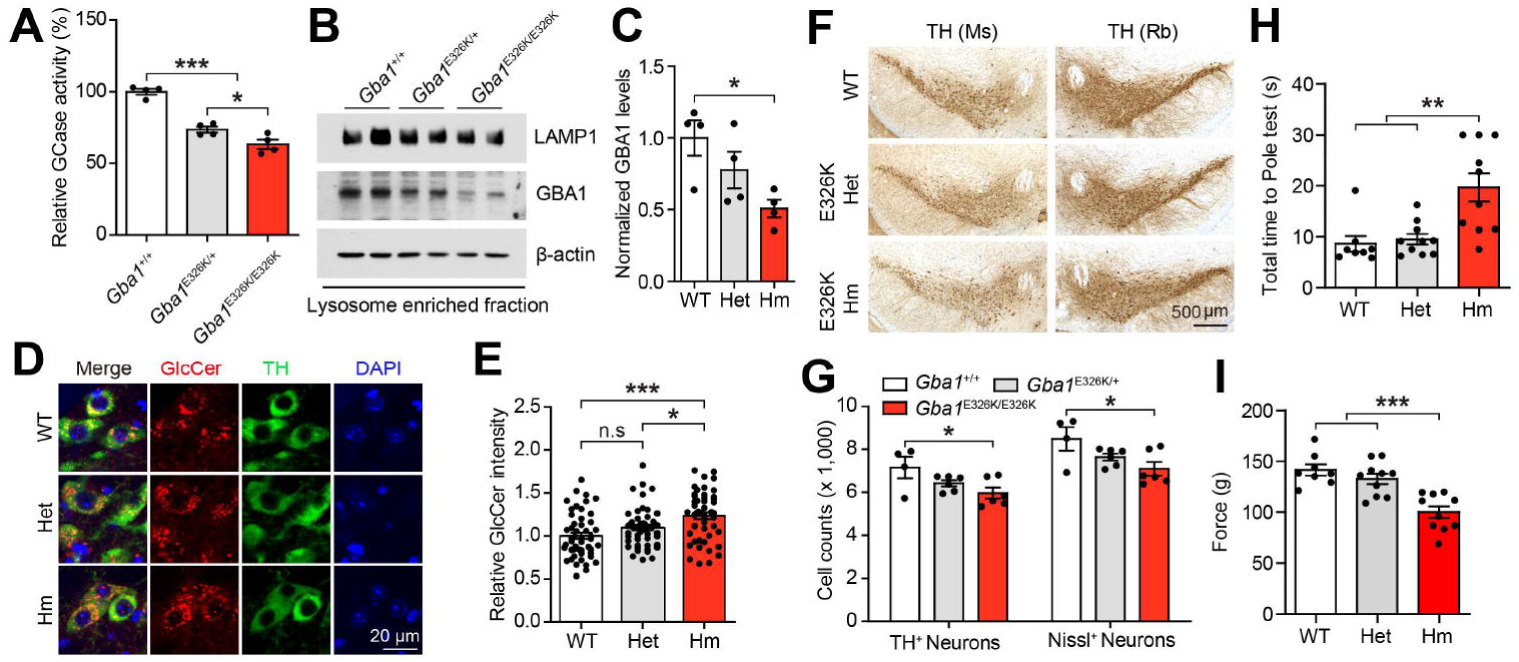
Characterization of *Gba1* E326K KI mice at 24 months of age. (A) GCase enzyme activity in the hippocampus of 24-month-old *Gba1* E326K KI mice including WT, Het, and Hm (n=4). (B) GBA1 protein levels were assessed using Western blot analysis in the lysosome-enriched fraction from the hippocampus. (C) Quantification of normalized GBA1 protein levels (n=4). (D) Representative immunostaining for GlcCer (red) and TH (green) in the SNc region. (E) Quantification of relative GlcCer intensity in TH-positive neurons (n=50). (F) Representative photomicrographs from coronal mesencephalon sections containing TH-positive neurons in the SNc region of 24-mont-old mice. (G) Stereology counts of the number of TH-positive and Nissl-positive neurons in the SNc region. Unbiased sterologic counting was performed in the SNc region (n=4-6). Results of mice on the (H) pole test, and (I) forelimb grip strength test (n=8-10). The error bars represent the S.E.M. \**P* < 0.05, \*\**P* < 0.01, \*\*\**P* < 0.001. n.s., not significant.

Immunofluorescence staining with glucosylceramide (GlcCer), a substrate of GCase, revealed an increase in the SNc of *Gba1*^E326K/E326K^ (Fig. 1D and E). Consistent with the findings in 24-month-old mice, we observed a significant reduction in both GCase activity and protein levels in the lysosome-enriched fraction of the ventral midbrain from 9-month-old *Gba1* E326K heterozygous and homozygous mice (Supplementary Fig. S2A-C). To further investigate the potential correlation between the reduced GCase enzyme activity and the loss of dopaminergic neurons in *Gba1* E326K mice, the number of tyrosine hydroxylase (TH)-positive neurons in the SNc was counted via an unbiased stereological analysis with different genotypes at 24 months of age (Fig. 1F and G). There was a significant loss of dopaminergic neurons in the SNc of *Gba1*^E326K/E326K^ at 24 months of age. However, there was no significant changes in the number of TH-positive neurons in *Gba1* E326K homozygous mice at 9 months of age (Supplementary Fig. S2D and E). Of note, the *Gba1* E326K homozygous mutation induced a mild accumulation of Triton X-100-insoluble α-syn (Supplementary Fig. S2F-H). Next, the motor function of the mice was assessed through pole test and grip strength test. *Gba1* E326K homozygous mice showed impaired performance in both tests at 24 months of age (Fig. 1H and I). However, no significant changes in motor function were observed at 9 months of age (Supplementary Fig. S2I and J). Collectively, the *Gba1* E326K mutation reduces the levels and activity of GCase in the lysosomes, ultimately resulting in the degeneration of dopaminergic neurons.

### Enhanced neuroinflammation and neuronal loss in the hippocampus of *Gba1* E326K KI mice

To assess the effect of the *Gba1* E326K mutation on neuroinflammation, we examined microglia and astrocytes by GFAP or Iba1 staining in the hippocampus and SNc of all genotypes. We observed a slight increase in microglia density in the SNc of *Gba1*^E326K/E326K^ mice at 9 months of age (Supplementary Fig. S3A and B). Furthermore, a significant increase in microglial density was observed in the dentate gyrus (DG) of the hippocampus of *Gba1*^E326K/E326K^ mice at 9 months of age (Supplementary Fig. S3C and D). However, no significant changes in microglial density were observed in the CA1 of the hippocampus of *Gba1*^E326K/E326K^ mice at 9 months of age (Supplementary Fig. S3C and E). The complement system plays critical roles in the central nervous system, as it is responsible for recognizing and eliminating invading pathogens as well as dead or modified self-cells, suggesting its significance in immune responses^27^. Complement activation has been found to be elevated in human PD brain, particularly in the presence of Lewy bodies (LB)^28^. To test whether the complement activation is evident in the presence of the E326K mutation, we performed Western blot analysis for C3 and Macrophage-Colony-Stimulating Factor Receptor (m-CSFR). The *Gba1* E326K mutation resulted in an increase of complement C3 protein levels. Additionally, the protein level of GFAP was markedly increased in the ventral midbrain of *Gba1* E326K homozygous KI mice at 9 months of age (Supplementary Fig. S3F and G). At 24 months of age, we did not observe a significant difference in GFAP immunoreactivity. However, we did observe a greater increase in Iba1 immunoreactivity in *Gba1* E326K homozygous KI mice compared to that in WT mice (Fig. 2A-C). Importantly, astrocytes and microglia were activated in the hippocampus of GBA1 E326K heterozygous and homozygous KI mice (Fig. 2D and E). In order to examine the effect of *Gba1* E326K mutation on neuronal cell death, an immunofluorescent labeling of NeuN was performed. Remarkably, the intensity of NeuN was decreased in the hippocampus of 24-month-old *Gba1* E326K heterozygous and homozygous mice compared to WT mice (Fig. 2F and G).

**Figure 2.**
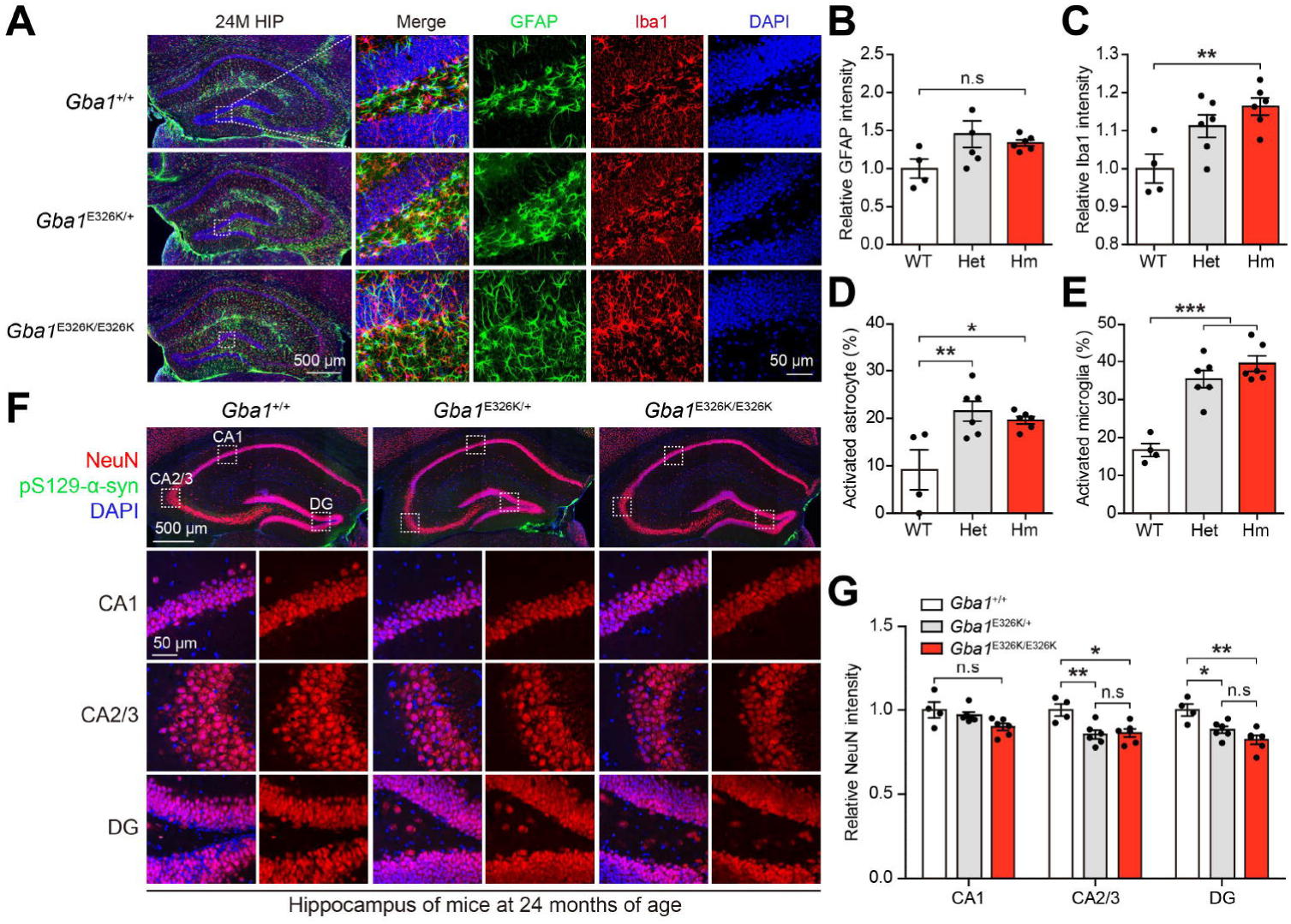
Neuroinflammation in the hippocampus of *Gba1* E326K KI mice. (A) Representative immunostaining images showing GFAP (green) and Iba1 (red) in the HIP region of 24-month-old mice. (B-C) Quantification of the relative intensity of (B) GFAP and (C) Iba1 in the dentate gyrus (n=4-6). (D-E) Percentage of activated (F) astrocyte and (G) microglia in the dentate gyrus (n=4-6). (F) Representative immunostaining images showing NeuN (red) and pS129-α-syn (green) in the HIP region of 24-month-old mice. (G) Quantification of the relative intensity of NeuN in CA1, CA2/3 and the dentate gyrus (n=4-6). The error bars represent the S.E.M. \**P* < 0.05, \*\**P* < 0.01, \*\*\**P* < 0.001. n.s., not significant.

### Exacerbated activation of microglia, formation of reactive astrocytes in α-syn PFF-induced *Gba1* E326K primary glia

A recent study has proposed that microglia activation can convert resting astrocytes to reactive astrocytes (A1 type) by secreting IL-1α, TNFα and C1q and that the conversion is toxic to neurons^29^. To determine whether α-syn PFF-treated microglia can induce reactive astrocytes, the expression of reactive astrocyte inducers was assessed in microglia after 3 h treatment with α-syn PFF. α-syn PFF treatment resulted in a significant increase in mRNA expression of *Tnfa, Il1a, Il1b, and Il6* in primary microglia as determined by quantitative PCR analysis (Fig. 3A-E). Notably, the *Gba1* E326K mutation conferred susceptibility to neuroinflammation triggered by α-syn PFF (Fig. 3A-E). Based on the observed neuronal loss and concurrent microglia activation associated with the *Gba1* E326K mutation, we hypothesized that the *Gba1* E326K mutation could exacerbate neuroinflammation in response to α-syn PFF-induced pathology. To determine the potential underlying mechanisms of neuroinflammation mediated by the *Gba1* E326K mutation, cultures of microglia, astrocytes, and neurons were assessed following treatment with α-syn PFF (Fig. 3A). Microglial-conditioned medium (MCM) derived from α-syn PFF-treated microglia was applied to astrocytes for 36 h. The resulting astrocyte-conditioned medium (ACM) was then collected, concentrated, and subsequently applied to mouse primary cortical neurons. Notably, the application of α-syn PFF-MCM preferentially induced markers of A1 astrocytes, and the expression of A1-specific transcripts were further increased in primary astrocytes from *Gba1* E326K homozygous mice (Fig. 3F). Consistent with the induction of A1-specific transcripts, the application of α-syn PFF-ACM reduced the cellular viability in primary neurons. The α-syn PFF-ACM derived from *Gba1* E326K homozygous astrocytes exhibited an even greater increase in cell death (Fig. 3G). Taken together, the conversion of astrocytes into reactive A1 astrocytes, induced by α-syn PFF, results in increased cell death, particularly in the presence of the *Gba1* E326K mutation.

**Figure 3.**
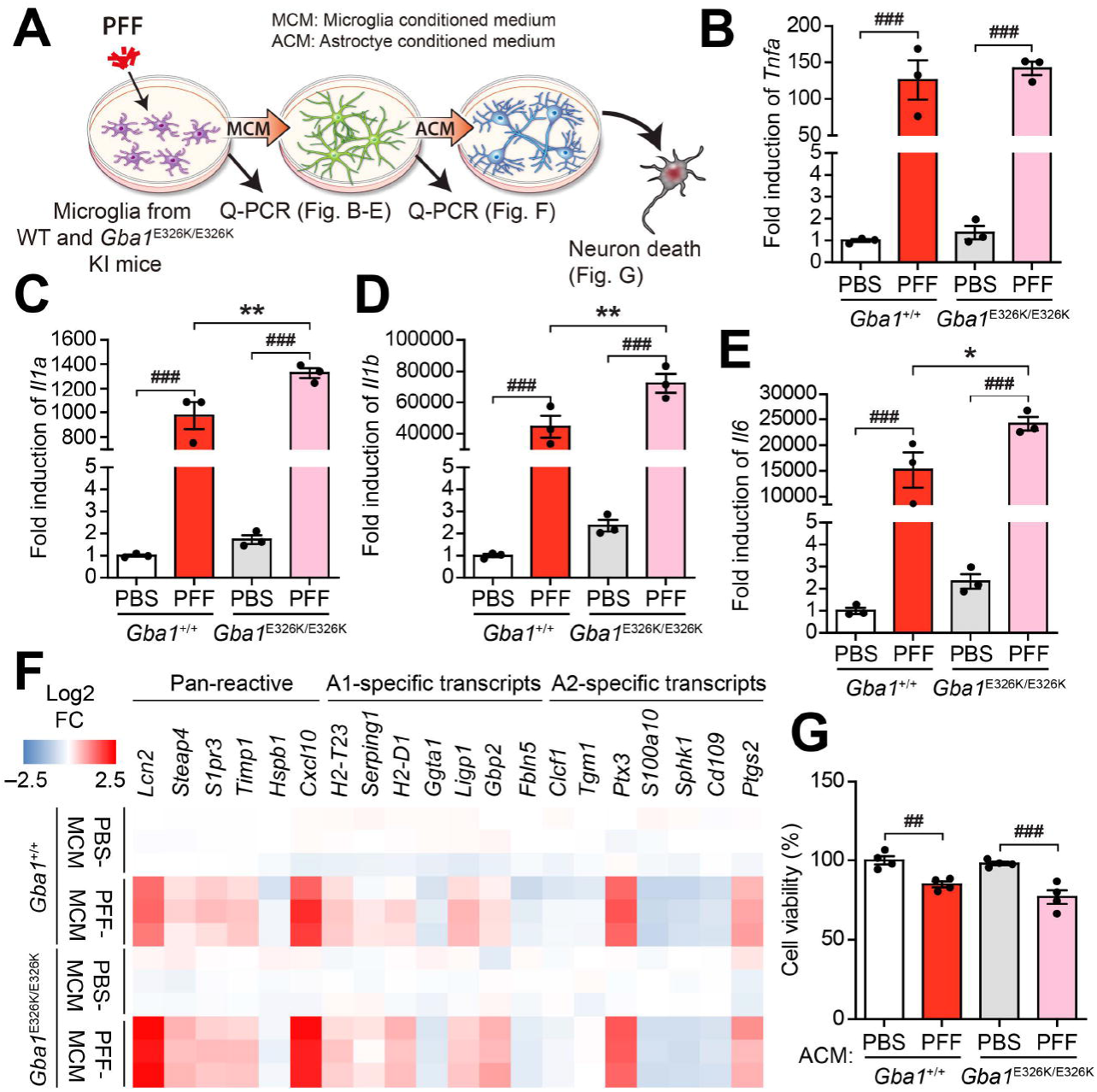
GBA1^E326K/E326K^ expression exacerbates microglia activation and reactive astrocytes formation due to PFF *in vitro.* (**A**) Experimental schematic diagram. (**B-E**) mRNA levels of reactive astrocytes inducers were measured in microglia 3h after PFF treatment using qPCR. (n=3). (**F**) Heatmap illustrating the qPCR results for levels of PAN-reactive, A1-specific, and A2-specific transcripts in primary astrocytes treated with microglia conditional medium (MCM) purified from PFF induced WT and GBA1^E326K/E326K^ primary microglia for 24 h (n=3). (**G**) Neuronal death was assessed using the AlamarBlue assay. Primary cortical neurons were treated with astrocytes conditioned medium (ACM) for 36 h. Error bars represent the mean ± S.E.M. WT vs Hm; \**P* < 0.05, \*\**P* < 0.01. PBS vs PFF; ^##^*P* < 0.01, ^###^*P* < 0.001

### Gut injection of **α**-Syn PFF induces exacerbated **α**-Syn pathology and neuroinflammation in the hippocampus of *Gba*1 E326K KI mice

Previous studies have suggested that pathologic α-syn has the ability to propagate from the gastrointestinal tract to the brain^24^. To test whether *Gba1* E326K KI mice have more progressive spreading of α-syn pathology from the gut, we utilized a mouse model of peripheral synucleinopathy by injecting mouse α-syn PFF into the duodenum of both WT and *Gba1* E326K KI mice. To monitor the progression of pathologic α-syn from the gut to the brain, immunostaining for Iba1 and pSer129-α-syn was performed 7 months after the injection of α-syn PFF. The intensity of pS129-α-syn in the hippocampus was increased by α-syn PFF. Notably, this increase was more pronounced in *Gba1* E326K KI mice (Fig. 4A and B). *Gba1* E326K homozygous KI mice showed enhanced microglia density even in the PBS-injected group, and the percentage of activated microglia was significantly higher in the hippocampus of the PFF-injected *Gba1* E326K homozygous KI mice 7 months post-injection (Fig. 4C and D). In *Gba1* E326K homozygous KI mice, Western blot analysis revealed a slight reduction in GBA1 protein levels and an increase of Triton X-100-soluble α-syn levels, regardless of PFF inoculation (Fig. 4E-G). Upon α-syn PFF injection into the gut, Triton X-100-insoluble α-syn and pSer129-α-syn accumulated in the hippocampus of mice. Remarkably, *Gba1* E326K KI mice exhibited an even higher accumulation of Triton X-100-insoluble α-syn and pSer129-α-syn in the hippocampus, following α-syn PFF injection (Fig. 4H-J). We also confirmed the presence of α-syn pathology and neuroinflammation in the amygdala 7 months after PFF injection. However, there were no notable alterations observed in the WT and KI groups (Supplementary Fig. S4A-D). Taken together, these results support that *Gba1* E326K KI mice exhibit enhanced α-syn pathology and neuroinflammation following the injection of α-syn PFF.

**Figure 4.**
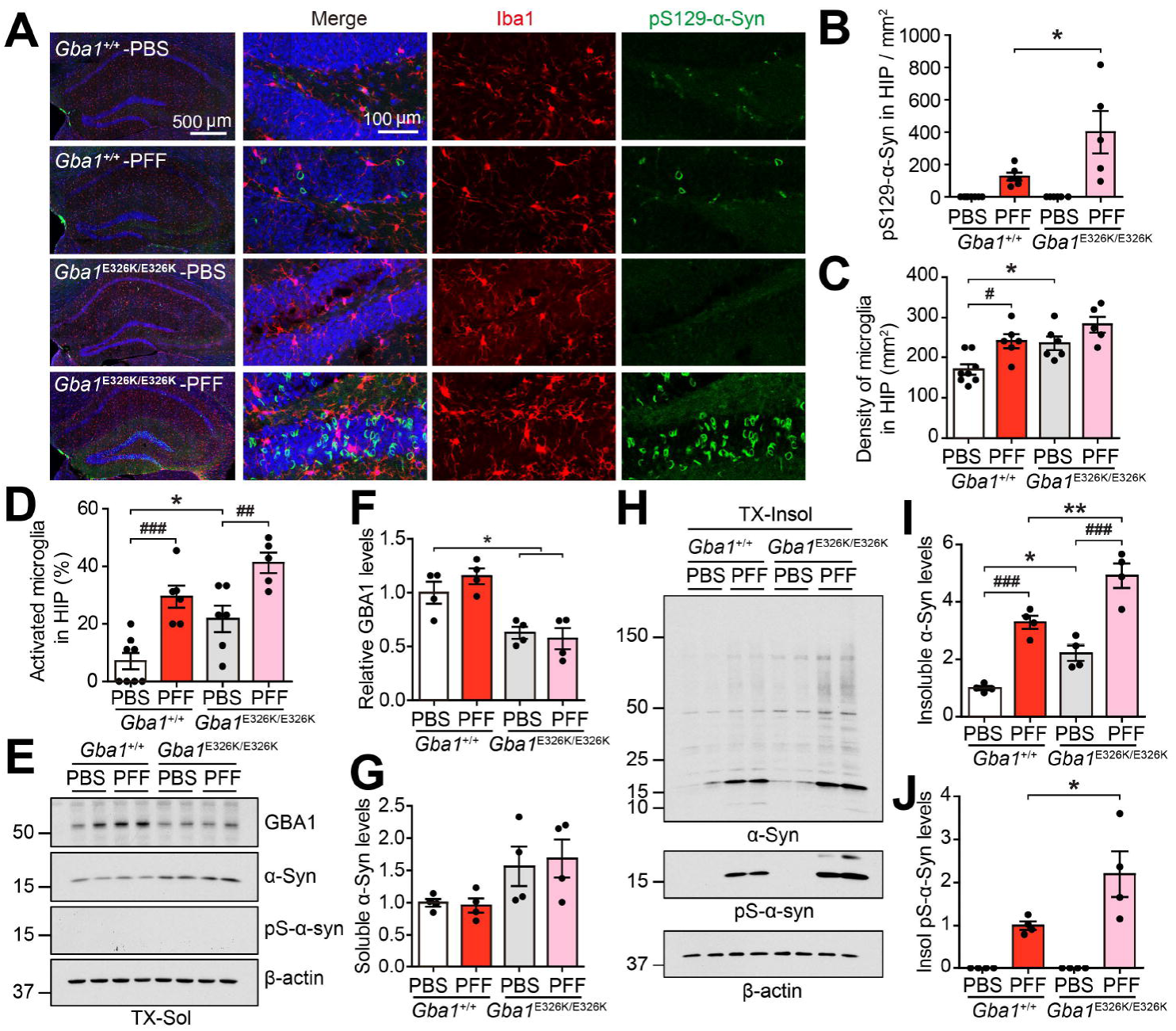
α-Syn pathology and neuroinflammation in the hippocampus of *Gba1* E326K KI mice induced by α-syn PFF injection into the gut. (A) Representative immunostaining images showing pS129-α-syn (green) and Iba1 (red) in the SNc region 7 months post-injection. (B) Quantification of the number of pS129-α-syn in the hippocampus. (C) Quantification of microglia density in the hippocampus. (D) Percentage of activated microglia in the hippocampus (n=5-8). (E) Representative immunoblots for GBA1, α-Syn and pS129-α-syn in the Triton X-100 soluble fraction. (F-G) Quantification of (G) GBA1 and (G) soluble α-Syn levels (n=4). (H) Representative immunoblots for α-Syn and pS129-α-syn in the Triton X-100 insoluble fraction. (I-J) Quantification of insoluble (E) α-Syn and (F) pS129-α-syn levels (n=4). Error bars represent the mean ± S.E.M. WT vs Hm; \**P* < 0.05, \*\**P* < 0.01. PBS vs PFF; **^##^***P* < 0.01, **^###^***P* < 0.001.

### Gut transmission of pathologic **α**-syn PFF exacerbates non-motor behavior deficits in *Gba1* E326K KI mice

Mice were subjected to the Morris water maze task 7 months PBS or α-syn PFF post-injection. On days 3 and 4 of exposure to the Morris water maze, the α-syn PFF-injected WT and *Gba1* E326K KI mice had a significantly prolonged escaped latency time compared to the PBS-treated mice. Notably, no significant difference in escaped latency time was observed between the WT and *Gba1* E326K KI mice (Fig. 5A). Following the last day of trial sessions (day 5), the α-syn PFF-injected WT and *Gba1* E326K KI mice had significantly decreased paths and swimming time in the target quadrant after the platform was removed. Notably, no significant difference in paths and swimming time in the target quadrant was observed between the WT and *Gba1* E326K KI mice. (Fig. 5B and C). There were no notable differences in swimming speed and total distance traveled among all the experimental groups (Supplementary Fig. S5A and B). Next, we conducted passive avoidance test to evaluate contextual fear conditioning, which is hippocampal-dependent or amygdala-dependent fear conditioning. The α-syn PFF-injected WT and *Gba1* E326K KI exhibited a lower step-through latency time compared to the mice injected with PBS. Notably, the α-syn PFF-injected *Gba1* E326K KI mice demonstrated a significantly reduced step-through latency time compared to the α-syn PFF-injected WT mice (Fig. 5D). Short-term or working memory was assessed using the spontaneous alternating behavior in the Y-maze test. The α-syn PFF-injected WT and *Gba1* E326K KI mice exhibited a significantly lower percentage of alternating behavior compared to the PBS-injected WT mice. Moreover, the α-syn PFF-injected *Gba1* E326K KI mice demonstrated even greater deficits in alternating behavior compared to the α-syn PFF-injected WT mice (Fig. 5E), No significant difference in the number of arm entries was observed among all the experimental groups (Supplementary Fig. S5C). Next, we performed a novel object recognition test to assess recognition memory^30^. The α-syn PFF-injected WT and *Gba1* E326K KI mice significantly reduced the recognition index and exploration time for novel objects compared to the PBS-treated mice. Notably, the α-syn PFF-injected *Gba1* E326K KI mice showed significantly greater deficits in recognition memory compared to the α-syn PFF-injected WT mice (Fig. 5F and G). The familiarization trial of exploration of the objects had no differences between WT and *Gba1* E326K KI (Fig. 5G). We subsequently performed several tests, including the open field test, elevated plus maze, forced swimming test, and tail suspension test, to assess neuropsychiatric features in the *Gba1* E326K homozygous KI mice. The open field test was used to assess movement patterns and symptoms of anxiety. The α-syn PFF-injected WT and *Gba1* E326K KI mice significantly reduced the time spent and the number of entries into the center zone compared to the PBS-treated mice, suggesting a change in basal anxiety levels (Fig. 5H-J). Importantly, the α-syn PFF-injected *Gba1* E326K KI mice showed a significantly greater reduction in the time spent and number of entries into the center zone compared to the α-syn PFF-injected WT mice (Fig. 5H-J). There were no notable differences in total distance moved and movement speed among all the experimental groups (Supplementary Fig. S5D and E). Anxiety was also measured by the elevated plus maze. On the elevated plus maze, the percentage of arm entries and time spent in the open arm was significantly decreased in the α-syn PFF-injected WT and *Gba1* E326K KI mice compared to the PBS-injected mice (Fig. 5K-M). Importantly, the α-syn PFF-injected *Gba1* E326K KI mice showed a significantly greater reduction in the percentage of arm entries and time spent in the open arm compared to the α-syn PFF-injected WT mice (Fig. 5K-M). To evaluate depressive-like symptoms, we performed a tail suspension test and forced swim test, the PFF-injected WT and PFF-injected *Gba1* E326K KI mice showed increased immobility times in both the forced swim test and tail suspension test compared to the PBS-injected mice (Fig. 5N and O). No significant difference was observed in the immobility times in the forced swim test between the α-syn PFF-injected *Gba1* WT and KI mice. However, in the tail suspension test, the PFF-injected *Gba1* E326K KI mice showed higher immobility times compared to the PFF-injected WT mice (Fig. 5N and O).

**Figure 5.**
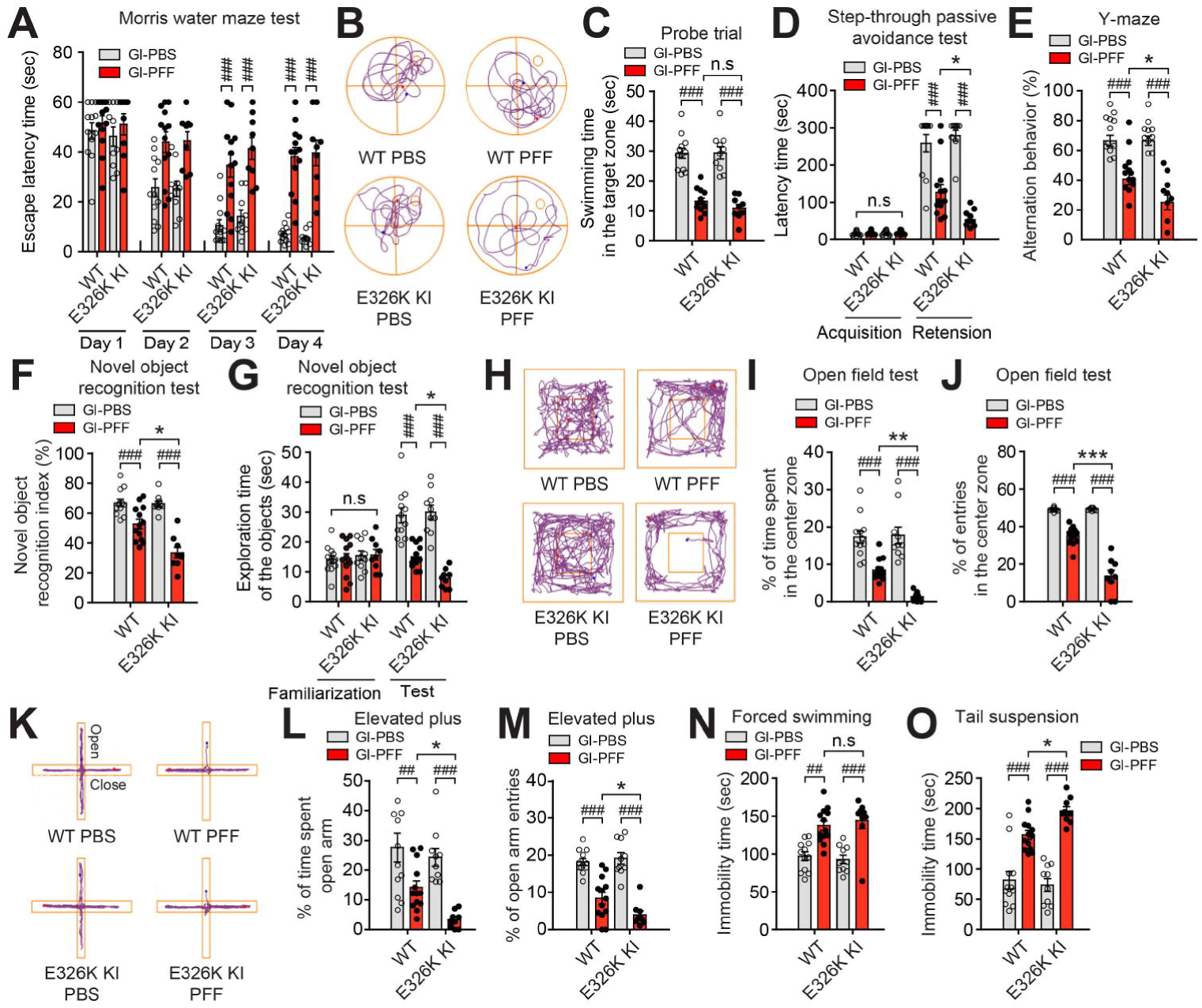
Non-motor symptoms in *Gba1* E326K KI mice induced by α-syn PFF injection into the gut. (A-C) Cognitive behavioral assessments 7 months after PBS and α-syn PFF gastrointestinal injection in WT and *Gba1* E326K KI homozygous mice (n=9-13). (A) Escape latency time. (B) Representative swimming paths of mice from each group in the Morris water maze test on the probe trial day 5. (C) Probe trial session in the Morris water maze test. The escape latencies were recorded (n=9-13). (D) Effect of *Gba1* E326K on learning and memory in the step-through passive avoidance test following α-syn PFF-induced hippocampal-dependent contextual or amygdala-dependent fear conditioning. For both training and test trials, the term of latency was the length of time it took a mouse to enter the dark compartment after the door opened (n=9-13). (E) Effect of *Gba1* E326K on α-syn PFF-induced short-term or working memory in the Y-maze test. Percentage of alternative behavior in the Y-maze test (n=9-13). (F) Percentage of novel object recognition index and (G) Exploration time of objects in the novel object recognition test (n=9-13). (H-J) Effect of *Gba1* E326K mutation on α-syn PFF-induced locomotion and central activity in the open-field test. (H) Representative movement paths of mice from each group in the open-field test. (I) The data of percentage of time spent and (J) entries in the center zone in the open-field test. (K) Representative movement paths of mice from each group in the elevated-plus maze. (L) Percentage of time spent in the open arm and (M) Open arm entries in the elevated-plus maze. (N-O) Effect of *Gba1* E326K mutation on α-syn PFF-induced depressive-like behavior in the (N) forced swim test and tail suspension test (O). During the final 4 min of a total 6 min test, the immobility times were recorded using the video tracking system (Any-Maze software). Error bars represent the mean ± S.E.M. WT vs Hm; \**P* < 0.05, \*\**P* < 0.01, \*\*\**P* < 0.001. PBS vs PFF; ^##^*P* < 0.01, ^###^*P* < 0.001. n.s., not significant.

## Discussion

Idiopathic PD is a progressive neurodegenerative disorder that manifests both motor and non-motor symptoms accompanying the accumulation of pathologic α-syn protein throughout multiple nervous systems, extending beyond the nigrostriatal system and even affecting the neocortex^31–34^. In addition to the motor symptoms, idiopathic PD patients suffer from cognitive dysfunctions, which encompass progressive cognitive decline, loss of attention, visual hallucinations, as well as anxiety and depression^35^. Mutations in the *GBA1* gene represent the strongest risk for developing sporadic PD and DLB^1,15^. Patients with *GBA1* mutations typically experience an earlier onset of the disease and exhibit more pronounced cognitive changes compared to those without *GBA1* mutations^17,36^. However, there is little information on how *GBA1* mutations affect cognitive symptoms to date.

The loss of catalytic activity and the reduction in protein levels of GCase within specific cellular compartments may contribute to the potential augmentation of disease progression. Notably, the activity of GCase in the lysosomal fraction may be a potential determinant of the severe aspects associated with the development of PD. In this study, the reduction of GCase protein levels in the *Gba1* E326K mutation resulted in the downregulation of GCase activity. Consequently, there was an induction of α-syn protein accumulation. Importantly, in vitro studies demonstrated that the *Gba1* E326K mutation significantly exacerbated α-syn PFF-induced neuronal toxicity and increase susceptibility to neuroinflammation, as well as the deposition of LB in the hippocampus of *Gba1* E326K KI mice. To understand the mechanism underlying the *Gba1* E326K mutation, we conducted a longitudinal collection of mouse sample. Our findings demonstrated that the presence of different phenotypes in aged mice, characterized by variations in GCase activity, function, and an increase in gliosis, which suggested that disease susceptibility as determined by activity and protein expression is highly dependent on this single mutation of the GCase protein. Consistent with this notion, heterozygous and homozygous *Gba1* E326K KI mice displayed a decrease in GCase activity, which was dependent on the gene dosage (Fig. 1). At 24-month-old mice, both GlcCer deposits in TH-positive neurons and dopaminergic neurodegeneration had progressed more significantly in homozygous *Gba1* E326K KI mice than in heterozygous *Gba1* E326K KI mice. These differences were correlated with variations in GCase activity. Furthermore, we utilized the gut-to-brain α-syn transmission mouse model to explore the effects of homozygous *Gba1* E326K KI mice. Our finding revealed an elevation in pathological α-syn aggregation and propagation from the gut to the brain in the presence of the *Gba1* E326K mutation. Consequently, we observed a significant increase in the phosphorylation of serine 129 of α-syn in brain lesion, accompanied by neuroinflammation and non-motor behavioral deficits.

Chronic inflammation, mediated by microglia, plays a pivotal role in the progressive degeneration of dopaminergic neurons in the brain of PD^37^. Recent studies suggested that activated microglia secrete TNFα, IL-1α, and C1q, which induce the conversion of resting astrocytes to reactive A1-type astrocytes, which exhibit distinct transcriptional profile^29,38,39^. Reactive A1 astrocyte releases unidentified neurotoxins, including complement factors, which contribute to the demise of neuronal cells^29,40^. Intriguingly, our findings revealed that the levels of TNFα, IL-1α, and complement C3, conversion of reactive astrocytes, and neuronal cell death induced by α-syn PFF were notably exacerbated in microglia expressing *Gba1* E326K (Fig. 3). Consistent with the in vitro findings, the α-syn PFF-injected Gba1 E326K KI mice showed a significant increase in the density and percentage of activated microglia in the hippocampus, along with a massive deposition of Lewy bodies (LB) when compared to α-syn PFF-injected wild-type (WT) mice (Fig. 4). Given that activated microglia have been reported to potentially promote α-syn aggregation and cell-to-cell transmission of pathologic α-syn^41^, it is worth to assess the primary role of microglia expressing Gba1 E326K in relation to LB deposition induced by α-syn PFF inoculation. However, it is important to note that this study only utilized mice after mating with ubiquitous Cre-expressing mice, such as CMV-Cre mice, due to the neonatal lethality of homozygous floxed mice. Therefore, the potential pathological role of the Gba1 mutation, whether it is cell-autonomous, non-cell-autonomous, or a combination of both, in contributing to LB deposition and behavioral symptoms, warrants further investigation. This would require future studies using neuron-specific or microglia-specific Cre expression mice to elucidate the precise mechanisms involved.

Taken together, our findings indicate that the E326K mutation-associated GBA1 deficiency enhances susceptibility to pathologic α-syn. Mutations in the GCase protein may confer a disadvantage in the survival of dopaminergic neurons, making them more susceptible to toxicity induced by α-syn PFF-induced both from the brain and the gut.

## Abbreviations

PD: Parkinson’s disease
PFF: Preformed fibrils
SNc: substantia nigra pars compacta
KI: Knock-in
GlcCer: glucosylceramide
CBE: Conduritol B epoxide
ACM: astrocyte-conditioned medium
MCM: microglia-conditioned medium

## Acknowledgment

We thank JHU transgenic core laboratory for assistance with the construction of heterozygous *Gba1* floxed mice by CRISPR service.

## IP rights notice

For the purpose of open access, the author has applied a CC-BY public copyright license to the Author Accepted Manuscript (AAM) version arising from this submission.

## Data availability statement

The authors affirm that data supporting the findings of this study included in this manuscript and additional supporting data are reasonably available from the corresponding author upon request.

## Supplementary material

Supplementary material is available at *BioRvix* online.

## References

1. Sidransky E, Nalls MA, Aasly JO, et al. Multicenter analysis of glucocerebrosidase mutations in Parkinson’s disease. N Engl J Med. 2009;361:1651–61.

2. Boer DEC, van Smeden J, Bouwstra JA, Aerts J. Glucocerebrosidase: Functions in and Beyond the Lysosome. J Clin Med. 2020;9:736

3. Bae EJ, Yang NY, Lee C, et al. Loss of glucocerebrosidase 1 activity causes lysosomal dysfunction and alpha-synuclein aggregation. Exp Mol Med. 2015;47:e188.

4. Kaye EM, Ullman MD, Wilson ER, Barranger JA. Type 2 and type 3 Gaucher disease: a morphological and biochemical study. Ann Neurol. 1986;20:223–30.

5. Chahine LM, Qiang J, Ashbridge E, et al. Clinical and biochemical differences in patients having Parkinson disease with vs without GBA mutations. JAMA Neurol. 2013;70:852–8.

6. Pandey MK, Burrow TA, Rani R, et al. Complement drives glucosylceramide accumulation and tissue inflammation in Gaucher disease. Nature. 2017;543:108–112.

7. Rocha EM, Smith GA, Park E, et al. Sustained Systemic Glucocerebrosidase Inhibition Induces Brain alpha-Synuclein Aggregation, Microglia and Complement C1q Activation in Mice. Antioxid Redox Signal. 2015;23:550–64.

8. Vitner EB, Salomon R, Farfel-Becker T, et al. RIPK3 as a potential therapeutic target for Gaucher’s disease. Nat Med. 2014;20:204–8.

9. Enquist IB, Lo Bianco C, Ooka A, et al. Murine models of acute neuronopathic Gaucher disease. Proc Natl Acad Sci U S A. 2007;104:17483–8.

10. Vitner EB, Farfel-Becker T, Eilam R, Biton I, Futerman AH. Contribution of brain inflammation to neuronal cell death in neuronopathic forms of Gaucher’s disease. Brain. 2012;135:1724–35.

11. Wong K, Sidransky E, Verma A, et al. Neuropathology provides clues to the pathophysiology of Gaucher disease. Mol Genet Metab. 2004;82:192–207.

12. Aflaki E, Stubblefield BK, McGlinchey RP, McMahon B, Ory DS, Sidransky E. A characterization of Gaucher iPS-derived astrocytes: Potential implications for Parkinson’s disease. Neurobiol Dis. 2020;134:104647.

13. Horowitz M, Pasmanik-Chor M, Ron I, Kolodny EH. The enigma of the E326K mutation in acid beta-glucocerebrosidase. Mol Genet Metab. 2011;104:35–8.

14. Duran R, Mencacci NE, Angeli AV, et al. The glucocerobrosidase E326K variant predisposes to Parkinson’s disease, but does not cause Gaucher’s disease. Mov Disord. 2013;28:232–236.

15. Nalls MA, Duran R, Lopez G, et al. A multicenter study of glucocerebrosidase mutations in dementia with Lewy bodies. JAMA Neurol. 2013;70:727–35.

16. Davis MY, Johnson CO, Leverenz JB, et al. Association of GBA Mutations and the E326K Polymorphism With Motor and Cognitive Progression in Parkinson Disease. JAMA Neurol. 2016;73:1217–1224.

17. Mata IF, Leverenz JB, Weintraub D, et al. GBA Variants are associated with a distinct pattern of cognitive deficits in Parkinson’s disease. Mov Disord. 2016;31:95–102.

18. McNeill A, Magalhaes J, Shen C, et al. Ambroxol improves lysosomal biochemistry in glucocerebrosidase mutation-linked Parkinson disease cells. Brain. 2014;137:1481–95.

19. Schondorf DC, Aureli M, McAllister FE, et al. iPSC-derived neurons from GBA1-associated Parkinson’s disease patients show autophagic defects and impaired calcium homeostasis. Nat Commun. 2014;5:4028.

20. Farfel-Becker T, Vitner EB, Kelly SL, et al. Neuronal accumulation of glucosylceramide in a mouse model of neuronopathic Gaucher disease leads to neurodegeneration. Hum Mol Genet. 2014;23:843–54.

21. Magalhaes J, Gegg ME, Migdalska-Richards A, Doherty MK, Whitfield PD, Schapira AH. Autophagic lysosome reformation dysfunction in glucocerebrosidase deficient cells: relevance to Parkinson disease. Hum Mol Genet. 2016;25:3432–3445.

22. Sanyal A, DeAndrade MP, Novis HS, et al. Lysosome and Inflammatory Defects in GBA1-Mutant Astrocytes Are Normalized by LRRK2 Inhibition. Mov Disord. 2020;35:760–773.

23. Kim S, Yun SP, Lee S, et al. GBA1 deficiency negatively affects physiological alpha-synuclein tetramers and related multimers. Proc Natl Acad Sci U S A. 2018;115:798–803.

24. Kim S, Kwon SH, Kam TI, et al. Transneuronal Propagation of Pathologic alpha-Synuclein from the Gut to the Brain Models Parkinson’s Disease. Neuron. 2019;103:627–641 e7.

25. Malini E, Grossi S, Deganuto M, et al. Functional analysis of 11 novel GBA alleles. Eur J Hum Genet. 2014;22:511–6.

26. Grace ME, Ashton-Prolla P, Pastores GM, Soni A, Desnick RJ. Non-pseudogene-derived complex acid beta-glucosidase mutations causing mild type 1 and severe type 2 gaucher disease. J Clin Invest. 1999;103:817–23.

27. Dalakas MC, Alexopoulos H, Spaeth PJ. Complement in neurological disorders and emerging complement-targeted therapeutics. Nat Rev Neurol. 2020;16:601–617.

28. Yamada T, McGeer PL, McGeer EG. Lewy bodies in Parkinson’s disease are recognized by antibodies to complement proteins. Acta Neuropathol. 1992;84:100–4.

29. Liddelow SA, Guttenplan KA, Clarke LE, et al. Neurotoxic reactive astrocytes are induced by activated microglia. Nature. 2017;541:481–487.

30. Bevins RA, Besheer J. Object recognition in rats and mice: a one-trial non-matching-to-sample learning task to study ‘recognition memory’. Nat Protoc. 2006;1:1306–11.

31. Bohnen NI, Muller M, Frey KA. Molecular Imaging and Updated Diagnostic Criteria in Lewy Body Dementias. Curr Neurol Neurosci Rep. 2017;17:73.

32. Goedert M, Jakes R, Spillantini MG. The Synucleinopathies: Twenty Years On. J Parkinsons Dis. 2017;7:S51–S69.

33. Peng C, Gathagan RJ, Lee VM. Distinct alpha-Synuclein strains and implications for heterogeneity among alpha-Synucleinopathies. Neurobiol Dis. 2018;109:209–218.

34. Wong YC, Krainc D. alpha-synuclein toxicity in neurodegeneration: mechanism and therapeutic strategies. Nat Med. 2017;23:1–13.

35. McKeith IG, Boeve BF, Dickson DW, et al. Diagnosis and management of dementia with Lewy bodies: Fourth consensus report of the DLB Consortium. Neurology. 2017;89:88–100.

36. Do J, McKinney C, Sharma P, Sidransky E. Glucocerebrosidase and its relevance to Parkinson disease. Mol Neurodegener. 2019;14:36.

37. Qian L, Flood PM. Microglial cells and Parkinson’s disease. Immunol Res. 2008;41:155–64.

38. Zamanian JL, Xu L, Foo LC, et al. Genomic analysis of reactive astrogliosis. J Neurosci. 2012;32:6391–410.

39. Escartin C, Galea E, Lakatos A, et al. Reactive astrocyte nomenclature, definitions, and future directions. Nat Neurosci. 2021;24:312–325.

40. Yun SP, Kam TI, Panicker N, et al. Block of A1 astrocyte conversion by microglia is neuroprotective in models of Parkinson’s disease. Nat Med. 2018;24:931–938.

41. George S, Rey NL, Tyson T, et al. Microglia affect alpha-synuclein cell-to-cell transfer in a mouse model of Parkinson’s disease. Mol Neurodegener. 2019;14:34.

