## Supplementary Figure 1 to 5 and Figure legends, Supplementary Table 1 for "Linking *Gba1* E326K mutation to microglia activation and mild age-dependent dopaminergic Neurodegeneration"

***BioRvix***
**Supplementary information**

### Linking *Gba1* E326K mutation to microglia activation and mild age-dependent dopaminergic Neurodegeneration

Sin Ho Kweon[1,2,*], Hye Guk Ryu[3,*], Hyeonwoo Park[3], Saebom Lee[1,2,4], Namshik Kim[1,2], Seung-Hwan Kwon[1,2], Shi-Xun Ma[1,2], Sangjune Kim[3,#], and Han Seok Ko[1,2,5,#]

[*,#]These authors contributed equally to this work.

**This file includes**

**Supplementary Figure 1 to 5 and Figure legends**

**Supplementary Table 1**

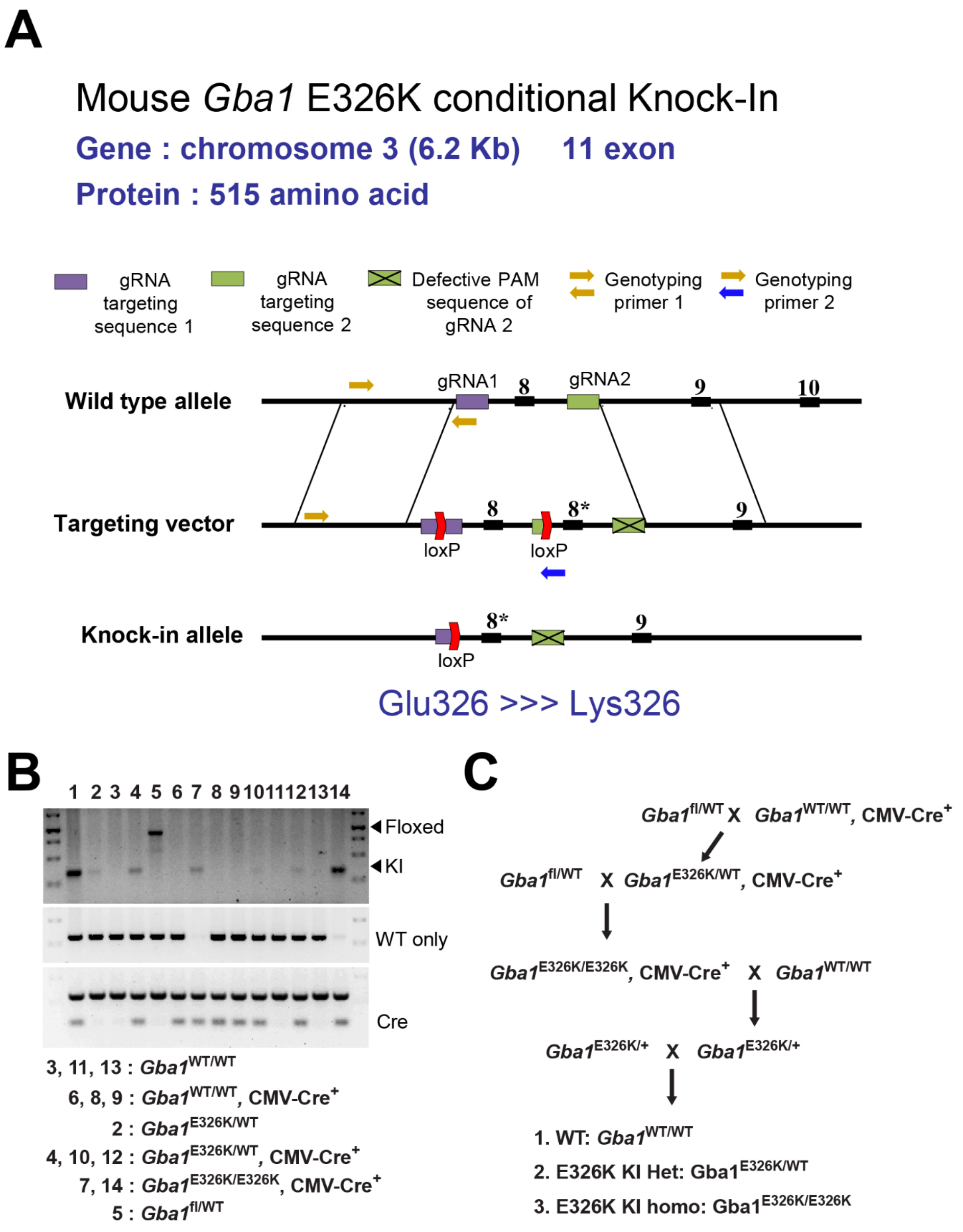


**Figure S1. Generation of E326K *Gba1* conditional KI mice.** *E326K Gba1* conditional KI mice were generated by homologous recombination using a targeting vector containing exon 8 flanked by loxP sites, followed by the mutated exon 8 (g1030G>A, Glu326Lys) and two sgRNA (1st sgRNA: 5'-gggccctggaaagttcagagtgg-3', 2nd sgRNA: 5'-tgtgaaagagaagataaccatgg-3'). Mice were then crossed with CMV-Cre transgenic mice to generate *E326K Gba1* mice. (A) Diagram illustrating the strategy for targeting E326K conditional KI mice. (B) Genotyping. (C) Breeding strategy.

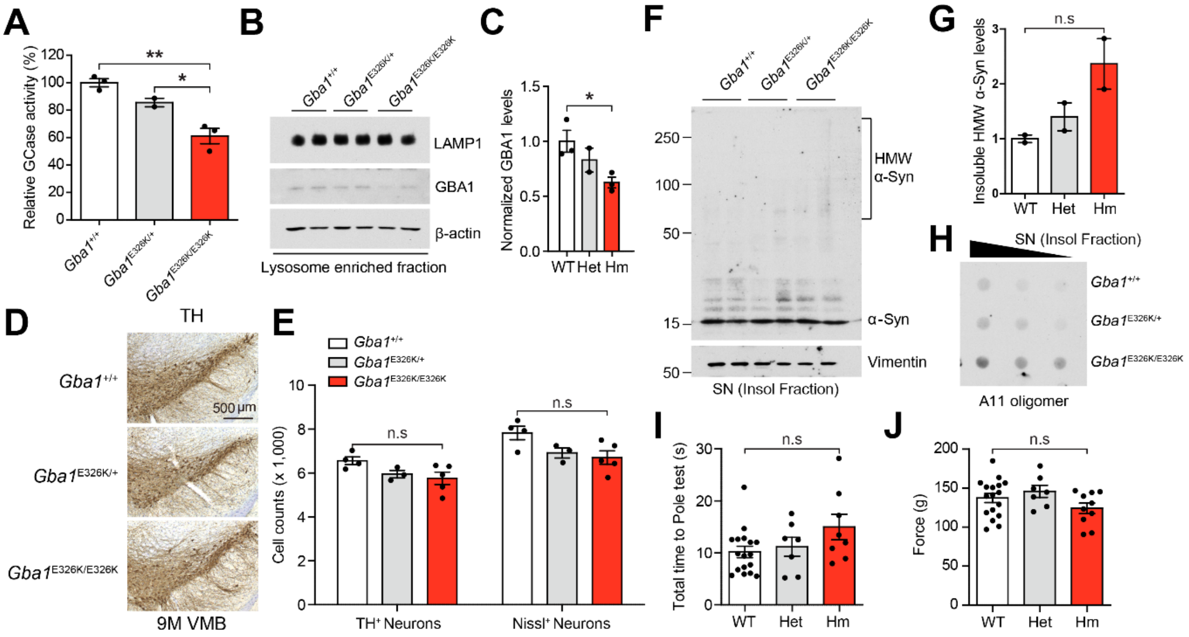


**Figure S2. Characterization of E326K *Gba1* KI mice**. (A) GBA1 enzyme activity in the ventral midbrain of 9-month-old WT, Het, and Hm of E326K *Gba1* KI mice (n=2-3). (B) GBA1 protein levels were assessed using Western blot analysis in the lysosome-enriched fraction from the ventral midbrain. (C) Quantification of normalized GBA1 protein levels (n=2-3). (D) Representative photomicrographs of coronal mesencephalon sections containing TH-positive neurons in the SNc region. (E) Stereology counts of TH- and Nissl-positive neurons in the SNc region. Unbiased stereologic counting was performed in the SNc region (n=3-5). (F) Representative immunoblot for α-Syn in the Triton X-100 insoluble fraction. (G) Quantification of insoluble HMW α-Syn levels (n=2). (H) Dot blot assay with anti-A11 oligomer antibody in the Triton X-100 insoluble fraction of the ventral midbrain. (I-J) Results of mice on the (I) pole test, and (J) forelimb grip strength test (n=7-17). The error bars represent the S.E.M. *$P < 0.05$, **$P < 0.01$. n.s., not significant.

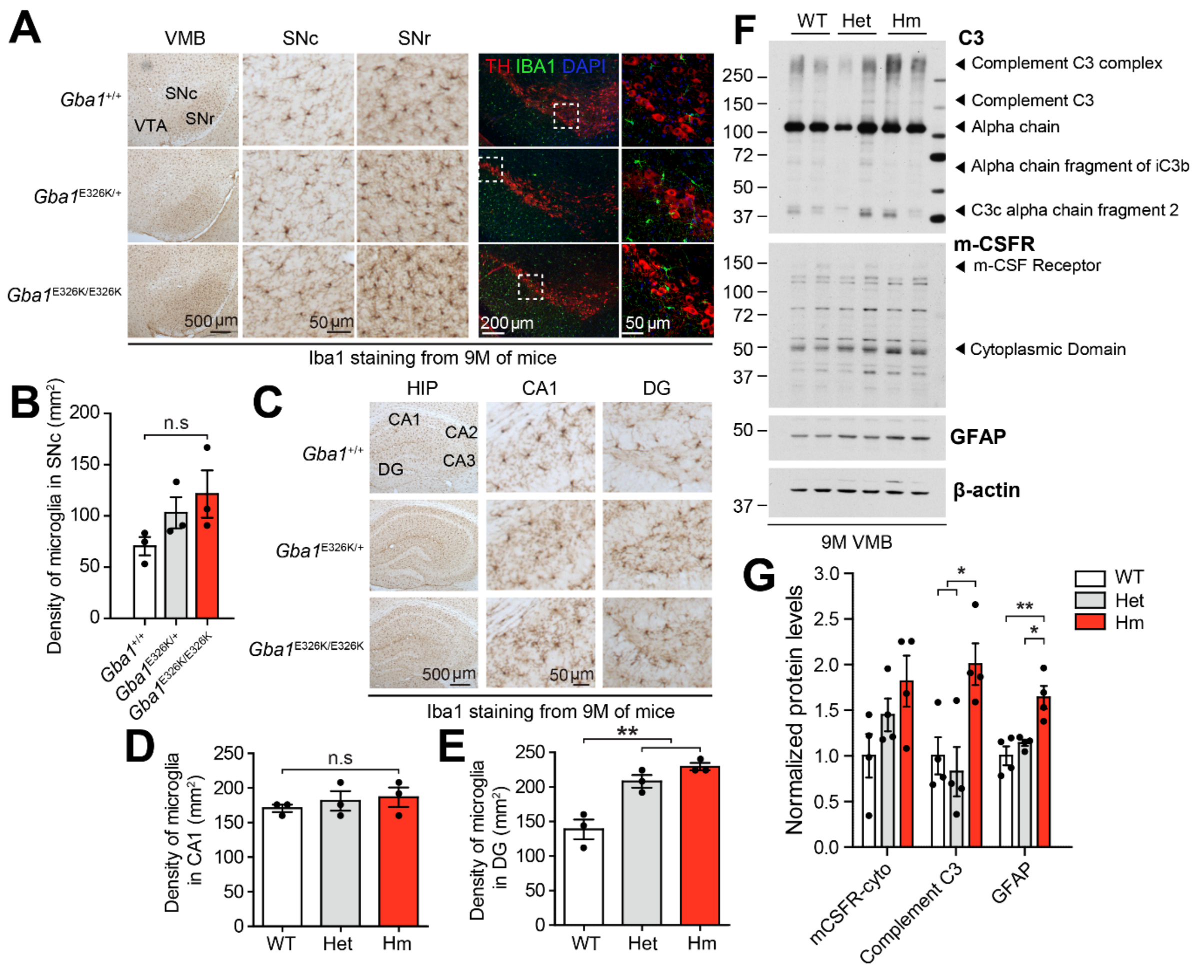


**Figure S3. Microglia activation in E326K *Gba1* KI mice at 9 months of age.** (A) Representative photomicrographs of coronal mesencephalon sections containing Iba1-positive microglia and immunostaining for Iba1 (Green) and TH (red) in the SNc region. (B) Quantification of microglia density in the SNc (n=3). (C) Representative photomicrographs of coronal mesencephalon sections containing Iba1-positive microglia in the hippocampus. (D) Density of microglia in the CA1 and (E) in the dentate gyrus (n=3). (F) Representative immunoblots for C3, m-CSFR and GFAP in the SNc region. (G) Quantification of normalized protein levels of m-CSFR-cytoplasmic domain, complement C3, and GFAP (n=4). The error bars represent the S.E.M. *$P$ < 0.05, **$P$ < 0.01. n.s., not significant.

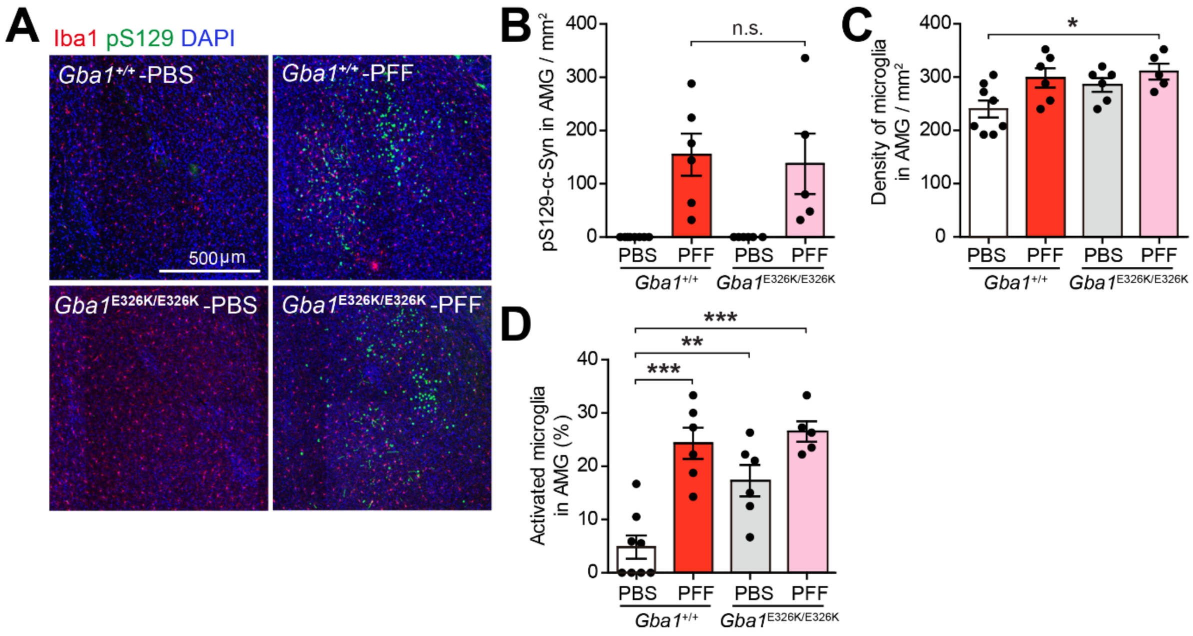


**Figure S4. α-Syn pathology and neuroinflammation in the amygdala of E326K *Gba1* KI mice induced by α-syn PFF injection into the gut.** (A) Representative immunostaining for pS129-α-syn (green) and Iba1 (red) in the amygdala region 7 months post-injection. (B) Quantification of the number of pS129-α-syn in the basolateral amygdala. (C) Quantification of the number of microglia in the basolateral amygdala. (D) Percentage of activated microglia in the basolateral amygdala (n=5-8). Error bars represent the mean ± S.E.M. *$P < 0.05$, **$P < 0.01$, ***$P < 0.001$. n.s., not significant.

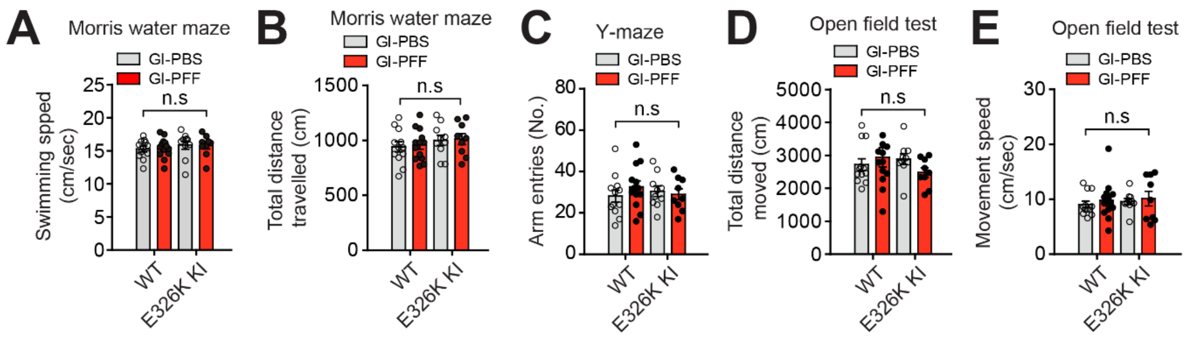


**Figure S5. Additional results for non-motor symptoms in E326K *Gba1* KI mice induced by α-syn PFF injection into the gut.** (A) Swimming speed and (B) total distanced traveled in probe trial sessions of the Morris water maze test (n=9-13). (C) Number of arm entries in the Y-maze test (n=9-13). The data of (D) total distance moved and (E) movement speed in the open-field test (n=9-13). Error bars represent the mean ± S.E.M.

**Table S1.** Primers used in this study

| Gene | Primer Sequence |
|---|---|
| *Tnfa* | F; CCCTCACACTCAGATCATCTTCT |
| | R; GCTACGACGTGGGCTACAG |
| *Il1a* | F; GCACCTTACACCTACCAGAGT |
| | R; AAACTTCTGCCTGACGAGCTT |
| *Il1b* | F; GCAACTGTTCCTGAACTCAACT |
| | R; ATCTTTTGGGGTCCGTCAACT |
| *Il6* | F; TAGTCCTTCCTACCCCAATTTCC |
| | R; TTGGTCCTTAGCCACTCCTTC |
| *Lcn2* | F; CCAGTTCGCCATGGTATTTT |
| | R; CACACTCACCACCCATTCAG |
| *Steap4* | F; CCCGAATCGTGTCTTTCCTA |
| | R; GGCCTGAGTAATGGTTGCAT |
| *S1pr3* | F; AAGCCTAGCGGGAGAGAAAC |
| | R; TCAGGGAACAATTGGGAGAG |
| *Timp1* | F; AGTGATTTCCCCGCCAACTC |
| | R; GGGGCCATCATGGTATCTGC |
| *Hspb1* | F; GACATGAGCAGTCGGATTGA |
| | R; GGATGGGGTGTAGGGGTACT |
| *Cxcl10* | F; CCCACGTGTTGAGATCATTG |
| | R; CACTGGGTAAAGGGGAGTGA |
| *H2-T23* | F; GGACCGCGAATGACATAGC |
| | R; GCACCTCAGGGTGACTTCAT |
| *Serping1* | F; ACAGCCCCCTCTGAATTCTT |
| | R; GGATGCTCTCCAAGTTGCTC |
| *H2-D1* | F; TCCGAGATTGTAAAGCGTGAAGA |
| | R; ACAGGGCAGTGCAGGGATAG |
| *Ggta1* | F; GTGAACAGCATGAGGGGTTT |
| | R; GTTTTGTTGCCTCTGGGTGT |
| *Ligp1* | F; GGGGCAATAGCTCATTGGTA |

| | |
|---|---|
| | R; ACCTCGAAGACATCCCCTTT |
| *Gbp2* | F; GGGGTCACTGTCTGACCACT |
| | R; GGGAAACCTGGGATGAGATT |
| *Fbln5* | F; CTTCAGATGCAAGCAACAA |
| | R; AGGCAGTGTCAGAGGCCTTA |
| *Clcf1* | F; CTTCAATCCTCCTCGACTGG |
| | R; TACGTCGGAGTTCAGCTGTG |
| *Tgm1* | F; CTGTTGGTCCCGTCCCAAA |
| | R; GGACCTTCCATTGTGCCTGG |
| *Ptx3* | F; AACAAGCTCTGTTGCCCATT |
| | R; TCCCAAATGGAACATTGGAT |
| *S100a10* | F; CCTCTGGCTGTGGACAAAAT |
| | R; CTGCTCACAAGAAGCAGTGG |
| *Sphk1* | F; GATGCATGAGGTGGTGAATG |
| | R; TGCTCGTACCCAGCATAGTG |
| *Cd109* | F; CACAGTCGGGAGCCCTAAAG |
| | R; GCAGCGATTTCGATGTCCAC |
| *Ptgs2* | F; GCTGTACAAGCAGTGGCAAA |
| | R; CCCCAAAGATAGCATCTGGA |
